# Signatures of introgression across the allele frequency spectrum

**DOI:** 10.1101/2020.07.06.189043

**Authors:** Simon H. Martin, William Amos

**Affiliations:** Institute of Evolutionary Biology, University of Edinburgh, Edinburgh, United Kingdom; Department of Zoology, University of Cambridge, Cambridge, United Kingdom

## Abstract

The detection of introgression from genomic data is transforming our view of species and the origins of adaptive variation. Among the most widely used approaches to detect introgression is the so-called ABBA BABA test or *D* statistic, which identifies excess allele sharing between non-sister taxa. Part of the appeal of *D* is its simplicity, but this also limits its informativeness, particularly about the timing and direction of introgression. Here we present a simple extension, *D* frequency spectrum or *D*_FS_, in which *D* is partitioned according to the frequencies of derived alleles. We use simulations over a large parameter space to show how *D*_FS_ caries information about various factors. In particular, recent introgression reliably leads to a peak in *D*_FS_ among low-frequency derived alleles, whereas violation of model assumptions can lead to a lack of signal at low-frequencies. We also reanalyse published empirical data from six different animal and plant taxa, and interpret the results in the light of our simulations, showing how *D*_FS_ provides novel insights. We currently see *D*_FS_ as a descriptive tool that will augment both simple and sophisticated tests for introgression, but in the future it may be usefully incorporated into probabilistic inference frameworks.

## INTRODUCTION

Hybridisation and introgression between related species occurs throughout the tree of life (Mallet et al. 2016). The continued growth of genome scale sequencing data now allows not only the detection of introgression, but also analysis of the fate of introgressed alleles. Such analyses provide insights into the timing of admixture events (Liu et al. 2014), the role of adaptive introgression (Racimo et al. 2017; Marburger et al. 2019) and the nature of species boundaries (Aeschbacher et al. 2017; Martin et al. 2019).

Among the most widely-used methods to detect intogression are the various variants of the ‘ABBA-BABA test’ *D* statistic (Green et al. 2010; Durand et al. 2011; Patterson et al. 2012). Given three ingroup populations and an outgroup with the relationship (((P1,P2),P3),O), *D* compares the number of derived alleles shared exclusively by P2 and P3 to that shared by P1 and P3. In the absence of introgression, such shared alleles restricted to non-sister taxon pairs can only emerge through incomplete lineage sorting (ILS) or recurrent mutation, and hence the two classes of sites are expected to be approximately equal in abundance (Green et al. 2010; Durand et al. 2011). An excess of one or the other is reflected in a non-zero *D,* and provides evidence for genetic exchange between P3 and either P1 or P2. The absolute value of *D* is determined by the amount of introgression as well as the amount of pre-existing shared variation due to ILS, and is therefore dependent on demography and population split times (Durand et al. 2011; Martin et al. 2015).

Although originally formulated for single sequences, *D* is equally applicable to samples of individuals by scaling according to the frequencies of derived alleles in each population (Durand et al. 2011). Even at sites where all populations are polymorphic, allele frequencies carry information about introgression, because shared ancestry causes frequencies to be correlated between populations (Patterson et al. 2012). This effect is also captured by the related f3 and f4 statistics (Patterson et al. 2012; Reich et al. 2012), which differ from *D* in that frequencies are not polarised by the use of an outgroup.

Although *D* provides a convenient measure of excess shared variation consistent with introgression, being a single number, it effectively averages over the entire allele frequency spectrum. By so doing, potentially valuable information about the history of the introgressed variants may be lost, including information that could help to distinguish true introgression from artefacts caused by violation of model assumptions. Specifically, ancestral population structure can result in an excess of shared ancestral polymorphisms between two non-sister taxa in the absence of introgression (Eriksson and Manica 2012). However, recent introgression and ancestral structure can be distinguished by considering the frequency distribution of shared derived alleles, as these should be more strongly biased toward lower frequencies in the case of introgression (Yang et al. 2012). Over time, anciently introgressed alleles can drift to higher frequencies and eventually become fixed, while others will be lost (Martin and Jiggins 2017). By averaging over all allele frequencies, *D* ignores information carried in the frequency distribution of introgressed alleles.

Here we introduce a simple descriptive measure that allows researchers to examine the nature of the signal underlying a non-zero *D*. The *D* frequency spectrum or *D*_FS_, reveals how the signal of introgression is broken down across different allele frequency classes (or bins). We use simulations to show that *D*_FS_ is strongly altered by different ages and directions of introgression, but can also be skewed by demographic events such as bottlenecks. In most cases, the signal of excess shared alleles is biased toward certain frequency bins, and may be entirely absent or even reversed at other frequencies. Even when there is no overall excess of shared variation (i.e. D~0), this may not be true across all frequency bins. We provide a tool that allows researchers to explore simulated *D*_FS_ over a large range of parameters. We then analyse published data from six plant and animal taxa and interpret the results in the light of our simulations. Overall, our findings show that *D*_FS_ provides additional information about the history of introgressed variation.

## NEW APPROACHES

*D*_FS_ is an extension to the ABBA BABA test *D* statistic (Figure 1). Both approaches aim to detect an excess of shared derived alleles between non-sister taxa, beyond those that are shared due to ILS alone. Whereas *D* averages across all sites in the genome, *D*_FS_ partitions the signal according to the frequency of the derived alleles in two focal populations, P1 and P2. Since shared derived alleles arising from both introgression and ILS may not be evenly distributed across allele frequencies, *D*_FS_ reveals how the signal of introgression (i.e. the excess in shared derived alleles) varies across allele frequency bins.

**Figure 1.**
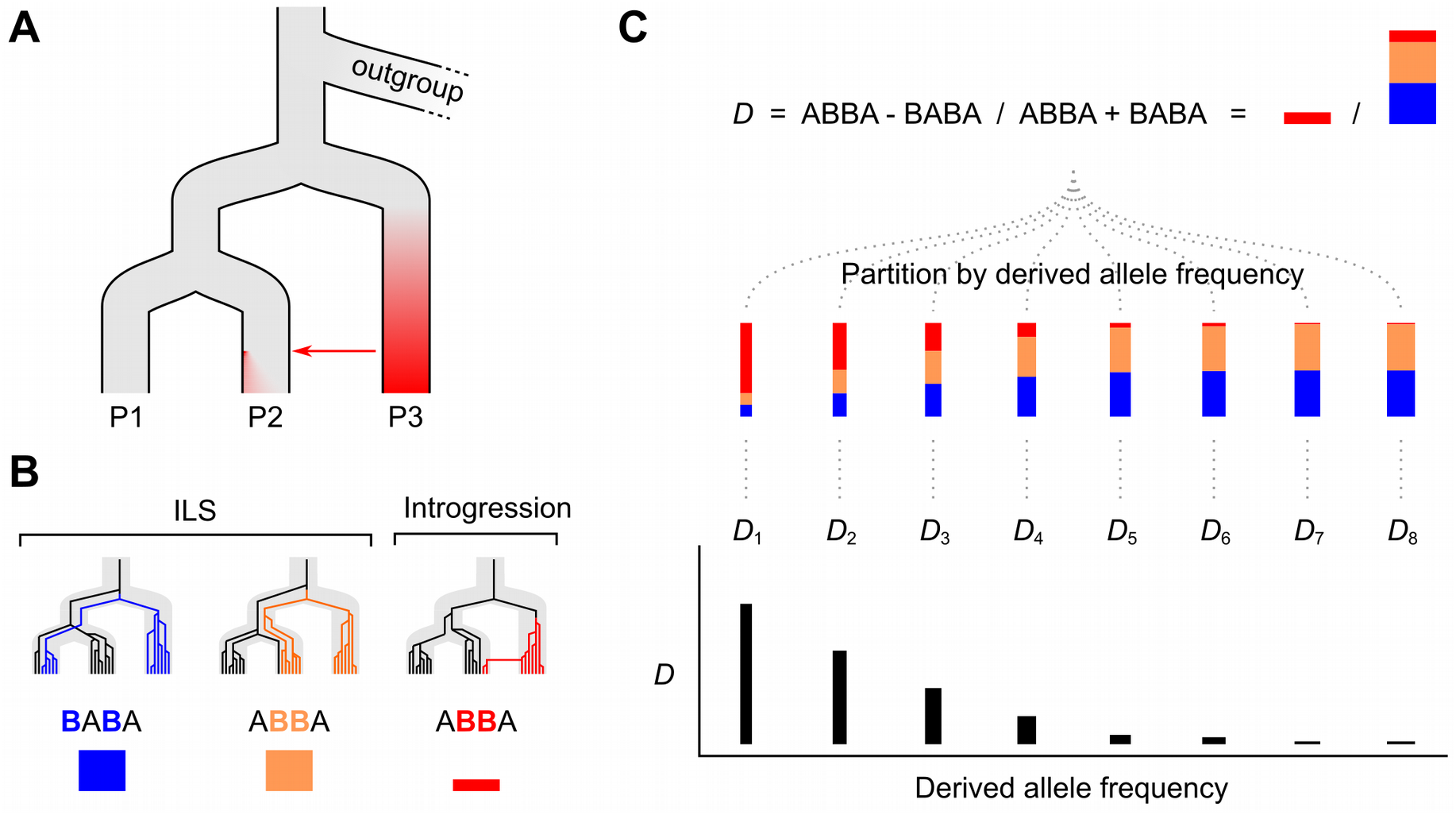
Conceptual representation of the *D* frequency spectrum. **A.** The approach makes use of three focal populations (P1, P2 and P3), in which gene flow is thought to have occurred between P3 and P2. The red shading represents the fact that divergent alleles will have arisen and spread within P3. When these introgress into P2, they will initially occur at low frequency. Through drift, some will increase in frequency over time and others will be lost, creating a distribution of frequencies of introgressed alleles. **B.** Three example genealogies that are all discordant with the population branching tree. The first two result from incomplete lineage sorting (ILS). Mutations on such genealogies lead to ‘BABA’ and ‘ABBA’ patterns in the sequence data, in which derived alleles (shown in colour) are shared between non-sister taxa. Because ILS involves deep coalescence, these shared derived alleles have time to drift, and can occur at any frequency in populations P1 and P2. The third genealogy represents introgression from P3 into P2. Because there is limited time for drift to occur, shared derived alleles resulting from recent introgression will tend to occur at lower frequency in the recipient population (P2). **C.** *D* is the difference in the observed number of ABBA and BABA patterns in the genome, normalised by their sum. *D* > 0 indicates an excess of ABBA due to introgression. *D*_FS_ is computed by partitioning ABBA and BABA counts according to the frequencies of derived alleles in both P1 and P2 (see Methods for details). With recent introgression, we expect *D*_FS_ to peak at low frequencies, because this is where ABBA patterns resulting from introgression will be most abundant relative to both the ABBA and BABA patterns resulting from ILS. The total number of sites contributing to each partition or ‘bin’ will differ, and each will contribute differently to the overall *D*. To show this, each bin is assigned a weighting, illustrated by the width of the vertical bars.

## RESULTS

### Simulation results

We used simulations of the site frequency spectrum over a broad range of parameters to explore how signatures of introgression in different allele frequency bins are affected by the timing, direction and rate of introgression, as well as by population sizes and split times. We provide an online tool where users can explore simulations covering over 68,000 parameter combinations: https://shmartin.shinyapps.io/shiny_plot_dfs_moments/. We also distribute code that can be used for exhaustive parameter exploration at https://github.com/simonhmartin/dfs. In the following results sections, we use representative simulations to demonstrate key features of *D*_FS_ under different evolutionary scenarios.

### Recent gene flow is most evident among low-frequency derived alleles

As expected, in simulations without gene flow, *D*_FS_ remains zero across all frequencies of the derived allele, provided population sizes remain constant. Simulating recent gene flow from P3 into P2 results in positive *D*_FS_ among bins representing low-frequency derived alleles (Figure 2A). When gene flow occurs further back in time, the signal of introgression tends to be more dispersed (Figure 2B), indicating that some introgressed alleles have drifted to higher frequencies. In the extreme case of very ancient gene flow, the signal becomes mainly restricted to the highest frequency bin (i.e. fixed derived alleles) (Figure 2C), indicating that all introgressed variation will eventually either go to fixation or be lost.

**Figure 2.**
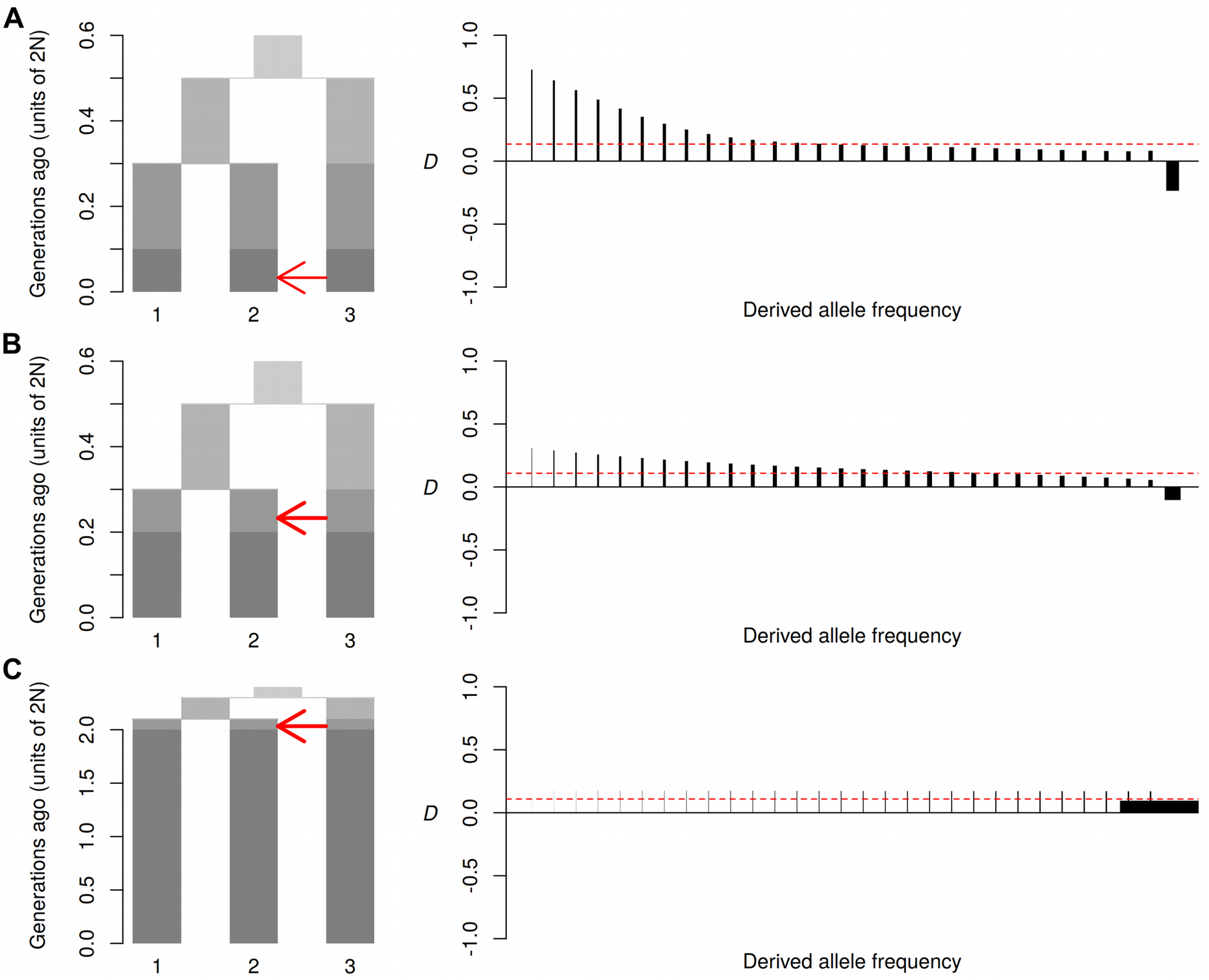
Effect of the timing of gene flow. Diagrams on the left show the simulated models of recent gene flow (A), older gene flow (B) and ancient gene flow (C). In each case, an ancestral population (top) splits into two daughter populations. One of these splits again to produce P1 and P2, and the remaining population becomes P3. The plot is divided into periods, shaded from light for the most ancient to dark for the most recent. Note the different scales of the y-axis. An arrow indicates a period in which gene flow occurs, and the direction of gene flow. Migration rate (2*Nm*) was set to 2 for recent gene flow (panel A) and 3 for both old (B) and very ancient (C) gene flow, to produce a comparable overall *D* value. Plots on the right show *D*_FS_. Each vertical line indicates the the stratified *D* value (*D*_k_) for each derived allele frequency bin. Widths of vertical lines are drawn in proportion to their weighting. Horizontal dashed red lines indicate the overall *D* value.

Unexpectedly, in simulations with recent gene flow from P3 into P2, the highest frequency bin often shows a negative *D* signal, reflecting an excess of derived alleles that are fixed in P1 and shared with P3, despite an overall positive *D* (Figure 2A). This counter-intuitive signal can be understood by recalling that, when population split times are relatively short, large numbers of shared derived alleles (‘ABBAs’ and ‘BABAs’) are generated by incomplete lineage sorting (ILS) of polymorphisms in the ancestral population. Many of these will have had enough time to drift to fixation in P1 and P2, but might still be segregating in P3. At such sites, introgression from P3 to P2 will act to reduce the number of derived alleles that are fixed in P2 and shared with P3. P1 is unaffected by introgression and therefore retains its fixed derived alleles, creating the imbalance in the highest-frequency bin. To test this logic, we decreased the effective population size of the donor population P3 to reduce segregating derived alleles due to ILS. As expected, this reduces and eventually abolishes the inverted signal, while also increasing the overall *D* value (Figure S1). This test also revealed another unexpected pattern: when P3 is very small, *D*_FS_ can have a rounded shape, with a dip toward the lowest frequency bins (Figure S1C). This occurs when a majority of introgressing derived alleles are already fixed in P3 such that they tend to occur at intermediate frequency in P2 immediately after introgressing.

### Demographic changes shift the frequencies of shared derived alleles

In the above scenarios, the sizes of the focal populations have been held constant. However, population bottlenecks impact allele frequency spectra (Watterson 1984), so likely also impact *D*_FS_, regardless of whether there has been inter-breeding. In a scenario with no gene flow and a bottleneck in P2, *D*_FS_ becomes negative in bins representing low and intermediate frequency derived alleles, gradually increasing and becoming strongly positive in the highest-frequency bin (Figure 3A). This reflects the way a bottleneck increases drift, thereby reducing the relative number of segregating derived alleles at low to intermediate frequencies in P2 and increasing the number of fixed or high-frequency derived alleles, as well as those that have been lost. Derived alleles segregating in P1 remain unaffected by the bottleneck. The negative *D* values at low and intermediate frequencies reflect those remaining segregating derived alleles in P1 that are shared with P3 due to ILS. Conversely, the positive *D* at high frequencies reflects the large number of derived alleles in P2 that are now fixed or nearly fixed, and shared with P3 due to ILS. Importantly, overall *D* remains zero despite the dramatic shifts in allele frequencies following demographic changes, as this does not change the sum total of derived alleles in P1 or P2 that are shared with P3 (Durand et al. 2011).

**Figure 3.**
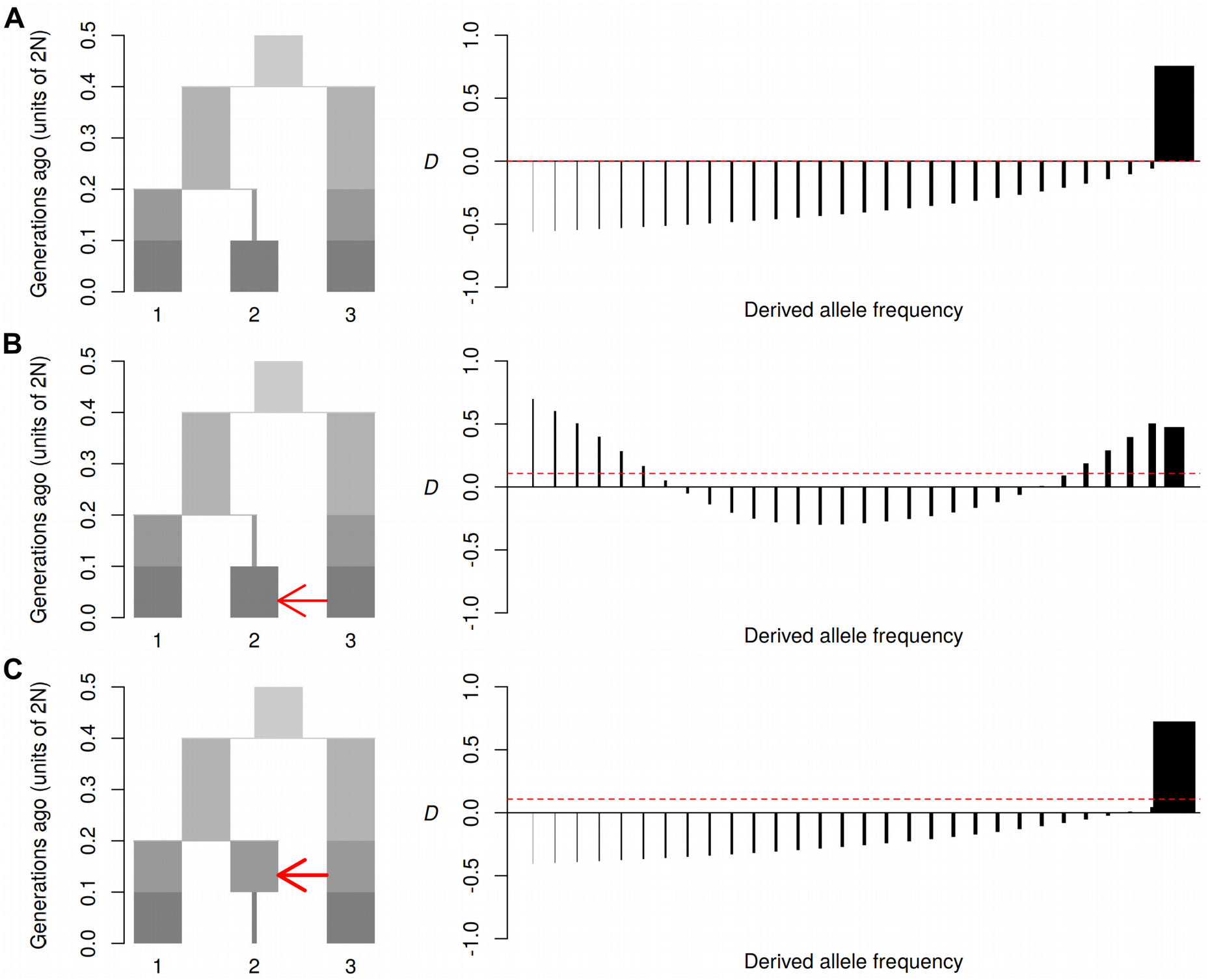
Effect of bottlenecks. Diagrams on the left show the simulated models of a bottleneck without gene flow (A), a bottleneck followed by gene flow (B) and a bottle neck after gene flow (C). Plots on the right show *D*_FS_ (see Figure 2 for details). Migration rate (2*Nm*) was set to 2 for recent gene flow (panel B) and 3 for older gene flow (C), to produce a comparable overall *D* value.

Introducing gene flow after the bottleneck shows that these two processes affect *D*_FS_ more or less additively (Figure 3B). There is a restoration of positive *D* values at low frequency due to introgression, whereas *D* values remain negative at intermediate frequencies, and positive at high frequencies due to the effects of drift described above. The overall *D* value becomes positive, confirming that *D* is able to capture the signal of introgression regardless of population size change (Durand et al. 2011). If gene flow occurs before the bottleneck, the positive *D*_FS_ at low frequencies is eliminated (Figure 3C). This is because introgressed alleles will tend either to be lost or to become fixed during the bottleneck, leaving few at low frequency.

### Behaviour under more complex scenarios

Above, we have considered the idealised case where there is unidirectional introgression from P3 into P2 and complete isolation of P1. We now examine the effect of relaxing these assumptions. Although *D* was first proposed under the assumption of introgression occurring ‘inward’ from P3 to P2, it is also able to detect intogression ‘outward’ from P2 to P3 (or from P1 to P3), albeit with reduced sensitivity (Martin et al. 2015). Our simulations show that gene flow outward from P2 to P3 generates a distinct *D*_FS_ signal that is approximately evenly dispersed across allele frequency bins (Figure 4A). This is unsurprising, because *D*_FS_ is stratified by derived allele frequencies in P1 and P2, but not P3. Any increase in shared derived alleles due to introgression from P2 into P3 should affect all allele frequency classes of the donor population (P2) approximately equally, regardless of the frequency distribution of introgressed alleles in the recipient population (P3). However, there is still variation in the weights of the different site classes, since those with lower frequencies of derived alleles contribute less to the overall *D* value. Adding bi-directional gene flow has an additive effect, and therefore restores the peak among low-frequency bins described above (Figure 4B). Importantly, bi-directional gene flow is detectable even if inward gene flow occurs at a much lower rate (e.g. ten fold lower) than outward gene flow (Figure S2).

**Figure 4.**
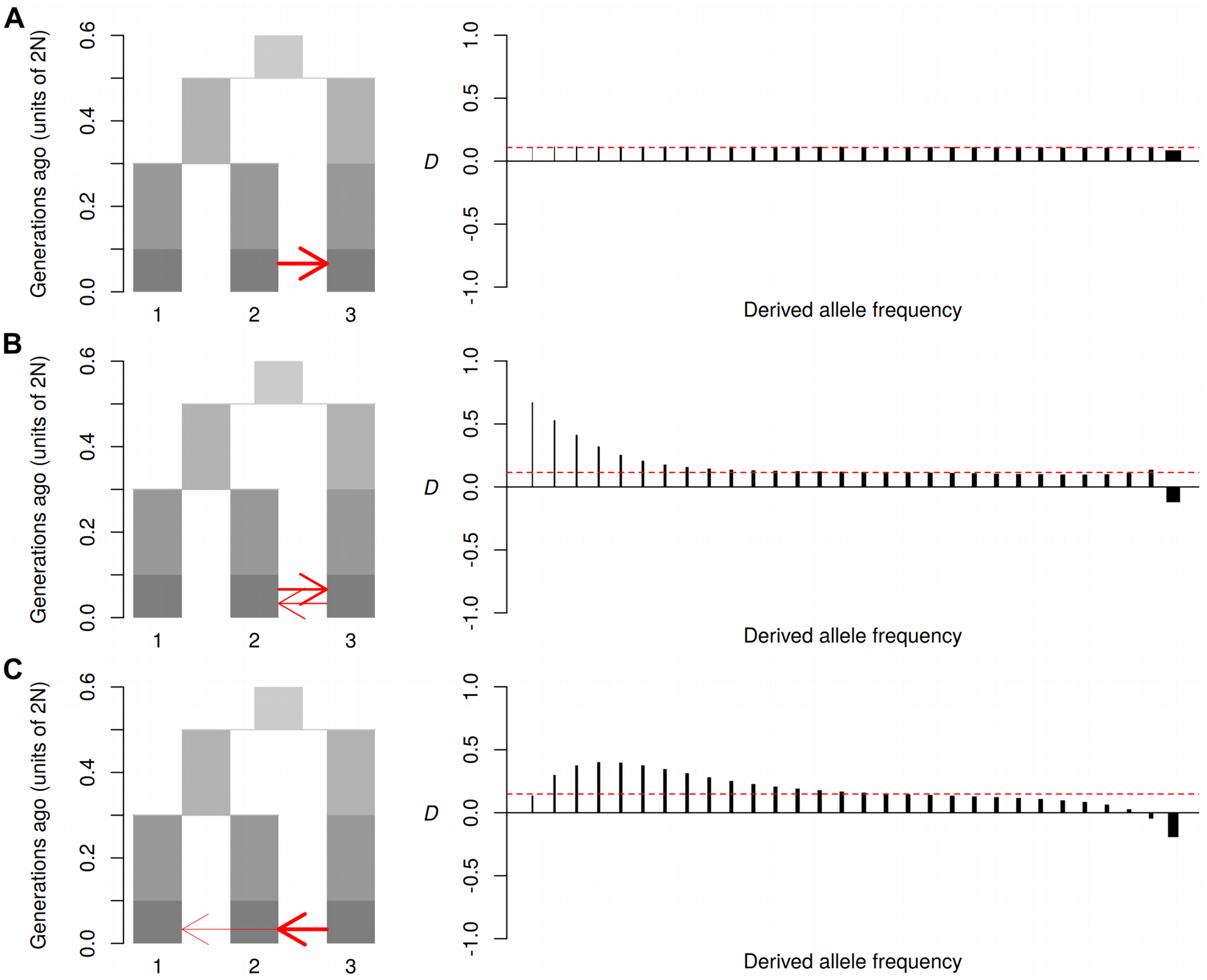
Behaviour under more complex scenarios. Diagrams on the left show the simulated models of “outward” gene flow from P2 to P3 (A)(2*Nm* = 2), bi-directional gene flow (B) (2*Nm* = 2 and 1 respectively) and gene flow from P3 into both P2 and P1 (C) (2*Nm* = 3 and 0.5, respectively). Plots on the right show *D*_FS_ (see Figure 2 for details).

Another scenario that is common in the real world is where populations P1 and P2 are not entirely isolated from one-another or from P3. Our simulations show that, provided introgression from P3 occurred recently, bi-directional gene flow between P1 and P2 has little impact on the shape of *D*_FS_, even if it occurs at a higher rate (Figure S3). However, unsurprisingly, if gene flow from P3 occurred deeper in the past, and bidirectional gene flow between P1 and P2 continued thereafter, it will eventually erode the signal of introgression (Figure S3).

If recent gene flow occurs from P3 into both P1 and P2, this can have a distinctive impact on the shape of *D*_FS_. If it occurs at the same rate into both P1 and P2, *D*_FS_ will of course become zero at all frequency classes, as there will be no excess of shared derived alleles. However, if gene flow from P3 to P1 occurs at a lower rate than from P3 to P2, this can produce a ‘rounding’ of the distribution, in which the signal is reduced in the lowest frequency bins, and peaks at a more intermediate frequency (Figure 4C). This again reflects the additive effect of gene flow on *D*_FS_. The low rate of gene flow from P3 to P1 causes these populations to share low-frequency derived alleles, thus offsetting the peak of low-frequency derived alleles shared between P2 and P3.

### Ancestral population structure

Finally, we consider the case of ancestral population structure introduced by Eriksson and Manica (2012) and Yang et al. (2012). We use a model similar to that of Yang et al., in which the ancestral population is structured into two sub-populations from before the divergence of P3, and this structure persists until the divergence of P1 and P2 (Figure S4). Thus, P2 and P3 will share an excess of ancestral variation as a result of the persistent population structure, despite the absence of recent gene flow. Our simulations show that this scenario leads to an unusual shape in *D*_FS_, in which the signal of excess sharing is restricted to intermediate and high-frequency derived alleles, and tends toward zero in the low-frequency bins (Figure S4). This finding is similar to that described by Yang et al., who used a related summary statistic (see Discussion).

### Empirical results

We applied *D*_FS_ to six published whole-genome data sets from *Heliconius* butterflies, tetraploid *Arabidopsis*, Ninespine sticklebacks (*Pungitius*), American sparrows (*Ammospiza*), North African date palms (*Phoenix*), and hominids. Based on previous work, we expected some of these taxa to show signatures of recent introgression, while others were expected to show signatures of longer term or more ancient introgression. In the case of humans, we expected the signal to also reflect the out-of-Africa bottleneck.

The North American Saltmarsh sparrow *Ammospiza caudacuta* and Nelson’s sparrow *A. nelsoni* are thought to have come into contact and begun hybridising only recently, after the last glacial retreat (Greenlaw 1993; Walsh et al. 2018). As expected, *D*_FS_ is skewed toward the lowest frequency variants and absolute values are low, consistent with a recent onset of inter-breeding (Figure 5A).

**Figure 5.**
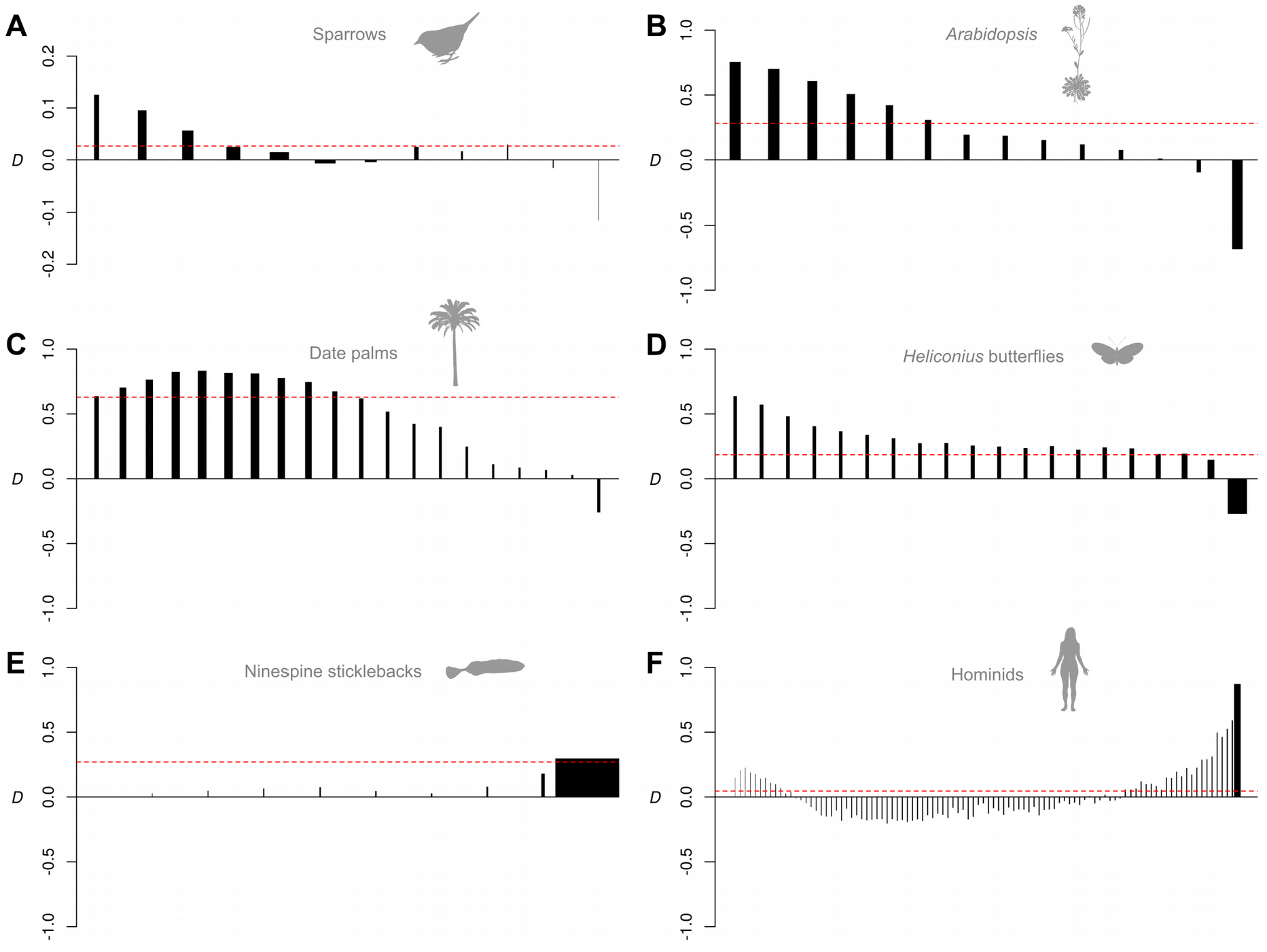
Analysis of six published data sets. *D*_FS_ for gene flow between sparrows *Ammospiza caudacuta* and *A. nelsoni* (A), tetraploid *Arabidopsis lyrata* and *A. arenosa* (B), Date palm *Phoenix dactylifera* and *P. theophrasti* (C), butterflies *Heliconius timareta* and *H. melpomene amaryllis* (D), sticklebakcs *Pungitius pungitius* and *P. sinensis* (E), and Neanderthals and humans (F). Each vertical line indicates the stratified *D* value (*D*_k_) for each derived allele frequency bin. Widths of vertical lines are drawn in proportion to their weighting. Horizontal dashed lines indicate the genome-wide *D* value. Differences in the number of bars among plots reflect the different sample sizes available for each system. Note that y-axis scales vary.

Tetraploid populations of *Arabidopsis lyrata* and *A. arenosa* are thought to have begun to hybridise extensively only after the emergence of tetraploid *A. lyrata*, around 80,000 generations ago (Marburger et al. 2019). *D*_FS_ is again skewed toward low frequencies but shows a much stronger signal than in the sparrows (Figure 5B), consistent with a higher rate of recent gene flow.

Date palm *Phoenix dactylifera* is thought to have experienced introgression from their wild Mediterranean relative *P. theophrasti* only after cultivation in North Africa (Flowers et al. 2019). *D*_FS_ indicates extensive gene flow, and has a rounded distribution that peaks at intermediate frequencies but declines toward low frequency bins (Figure 5C). This rounded distribution might reflect the small population size of the donor, *P. theophrasti* from Crete, which would cause more introgressed alleles to occur at intermediate frequency (e.g. Figure S1C). Alternatively a similar pattern could emerge if additional but less extensive gene flow has occurred from *P. theophrasti* into middle-eastern date palm (e.g. Figure 4C).

The butterflies *Heliconius timareta* and *H. melpomene amaryllis* are thought to have experienced long-term gene flow over millions of generations (Martin et al. 2013). *D*_FS_ has a bias toward low frequencies but also a broad tail in the higher frequencies (Figure 5D), consistent with longer term and/or bidirectional gene flow (see above). In another pair, *H. cydno* and *H. melpomene rosina*, the *D*_FS_ signal lacks the low-frequency peak (Figure S5B), suggesting that gene flow may be constrained to further in the past. This might reflect an increasing strength in the reproductive barrier due to processes such as reinforcement, for which there is experimental evidence in this pair (Kronforst et al. 2007). Although several other combinations of populations could be analysed in this *Heliconius* data set, few include a P1 population that is allopatric from P3 or a related species (Martin et al. 2019). Using P1 populations that are themselves partly admixed with P3 leads to more rounded peaks (Figure S5C, S5D) as expected based on our simulations above (Figure 4C).

In *Pungitius* sticklebacks, previous work has shown that an entire chromosome introgressed from *P. sinensis* into the ninespine stickleback *P. pungitius* to form a neo-sex chromosome, but that genome-wide introgression between the species has also occurred on autosomes (Dixon et al. 2019). *D*_FS_ for introgressed alleles in *P. sinensis* (exluding the neo-sex chromosome) shows a dramatic skew, with almost all shared derived alleles being fixed in the recipient population (Figure 5E). This is consistent with a deep age of introgression relative to the effective population size, such that most introgressed alleles have since drifted to fixation.

In humans, we expected the frequency of derived alleles shared between Europeans and Neanderthals to be shaped by both the out-of-Africa bottleneck and by recent introgression. *D*_FS_ is largely consistent with these expectations, with negative values at intermediate frequencies, reflecting the loss of segregating ancestral variants through drift during the bottleneck (see above). *D*_FS_ becomes positive at low-frequencies, consistent with retention of some introgressed variation at low frequency. The large sample size enables us to detect a weak decrease in the lowest frequency bin. Although our simulations suggest that this could be explained by limited introgression of Neanderthal alleles into African populations (also see Chen et al., 2020), we suggest that future studies interrogate this signal further.

## DISCUSSION

The detection of introgression using genomic data is transforming our view of species and the origins of adaptive variation. Various methods exist to make inferences about demographic history and introgression using rich summaries of genomic data such as the site frequency spectrum (Gutenkunst et al. 2009; Excoffier et al. 2013), patterns of linkage disequilibrium (Machado et al. 2002; Sankararaman et al. 2012) admixture tract lengths (Harris and Nielsen 2013) or combined signals (Lohse et al. 2016; Roux et al. 2016). Nevertheless, the ABBA BABA test retains widespread popular appeal due to its relative simplicity: it captures the key information relevant to the question of whether introgression has occurred in a single value. However, in doing so, *D* and its derivatives fail to capture information that could help better to interpret the signal. Allele frequencies carry additional information about how introgressed alleles are distributed in the recipient population, which reflects both the timing and quantity of introgression, along with other processes.

*D*_FS_ is an intuitive extension of *D* that greatly enhances its information content with minimal extra computation. Most importantly, the extent to which *D*_FS_ is biased toward low frequencies is a good indicator of the recency of introgression. Indeed, our simulations demonstrate that the low-frequency bins are the most sensitive for detection of recent introgression. The reason for this is two-fold. Firstly, under low to moderate rates of gene flow, introgressed alleles will tend to occur at low frequency in the recipient population as they have not had time to drift to higher frequency. An exception may be strongly positively-selected introgressed alleles, but these will typically represent a small proportion of the genome. Secondly, *D* captures the difference in numbers of sites carrying shared derived alleles (i.e. ABBAs and BABAs) normalised by the total number of ABBAs and BABAs. Because the majority of these will typically have arisen through incomplete lineage sorting, they will tend to involve derived alleles spread more or less uniformly across the entire frequency spectrum, and after sufficient time for genetic drift, all will involve only fixed derived alleles. Consequently, ABBAs and BABAs generated by incomplete lineage sorting tend to be rare in the lowest frequency bins, making these the most sensitive bins for detecting introgression.

A lack of signal of introgression among low-frequency derived alleles only emerges if introgression is very ancient or if the underlying scenario is more complicated, such as including additional introgression into the ‘control’ population (P1), or even no introgression at all but rather ancestral population structure. Unlike gene flow, the latter scenario results in excess sharing of high-frequency alleles that existed as polymorphisms in the ancestral population. Therefore, detection of low frequency shared derived alleles serves as a useful rule-of-thumb indicator when assessing the validity of claims of recent introgression. This signature was previously demonstrated by Yang et al. (2012) using a related summary called the ‘doubly conditioned frequency spectrum’ (*dcfs*). The *dcfs* also examines the frequency distribution of shared derived alleles, but unlike *D*_FS_ it does not explicitly test for an excess of shared derived alleles, and therefore cannot as easily distinguish between introgression and other signatures that could also skew allele frequencies.

Most of our analyses of empirical data show patterns consistent with recent introgression. The most extreme skew towards an excess of low-frequency shared derived alleles is seen in the saltmarsh and Nelson’s sparrows, which are thought to have come into contact only since the last glacial retreat (Greenlaw 1993; Walsh et al. 2018). At the opposite end of the spectrum are the Amur and ninespine sticklebacks, in which shared derived are almost entirely fixed in the Amur stickelback, implying that introgression was ancient relative to its effective population size. At present, although we have been able to identify a number of features of *D*_FS_ that can be linked to particular scenarios, the overall interpretation remains qualitative. However, we envisage that *D*_FS_ could in the future facilitate quantitative inferences of the extent and timing of gene flow by, for example, incorporation into inferential frameworks such as approximate bayesian computation (ABC) (e.g. Roux et al., 2016). Unlike full joint site frequency spectra, which can have vast numbers of entries with large sample sizes, *D*_FS_ retains relatively low dimensionality while also retaining high information content.

Demographic change is an important additional complicating factor for interpreting *D*_FS_. We explored this in the context of a severe bottleneck in one of the two focal populations. In the absence of gene flow, strong drift in the bottlenecked population leads to a distinctive pattern in which there is an excess of high-frequency or fixed derived alleles shared between the bottlenecked population and P3, and an opposite excess of derived alleles at low and intermediate frequencies shared between the non-bottlenecked population and P3. Gene flow after the bottleneck tends to have an additive effect on *D*_FS_, so in many cases it should still be possible to infer the presence of recent introgression in a bottlenecked population from an excess signal at low frequencies. On the other hand, if detection of bottlenecks is of interest, our findings reveal that considering the joint frequencies in multiple populations provides a sensitive indicator of population size change. It seems likely that analysis of joint frequency spectra could provide additional power to infer population size change over and above classical single-population methods (e.g. Liu and Fu, 2015). Nonetheless, it is important to remember that because both *D* and *D*_FS_ depend on ratios between populations, their values are dependent on the assumption of constant mutation rate which may, in some cases, be incorrect (Amos 2013; Mallick et al. 2016; Xie et al. 2016).

In conclusion, we recommend that researchers making use of both simple (e.g. the ABBA BABA test) and inference-based (e.g. maximum likelihood or ABC inference) approaches to investigate introgression, also include analysis of *D*_FS_ as part of their work-flow. We do not propose *D*_FS_ as a replacement for inference methods but more as an addition that allows further exploration of the genetic information space. This may be particularly valuable for exploratory studies in which little is known about direction and timing of introgression. Regardless of which methods of inference are used, it is important to understand the nature of the signal that leads to a particular inference. *D*_FS_ provides an intuitive descriptor that falls between the simple and sophisticated approaches, but retains advantages of both.

## METHODS

### The *D* frequency spectrum

We first define the *D* frequency spectrum (*D*_FS_) by relating it to the conventional *D* statistic. Given a sequence alignment of *l* sites, including sequences from three populations and an outgroup with the relationship (((P1, P2), P3), O), and assuming for now that we have just a single haploid sequence representing each taxon,

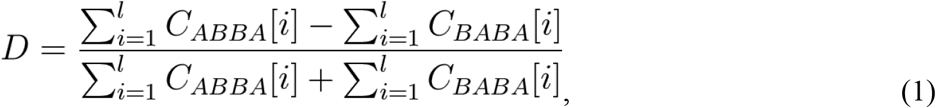

where *C*_*ABBA*_[*i*] and *C*_*BABA*_[*i*] are either 1 or 0 depending on whether the alignment at site *i* matches the ‘ABBA’ or ‘BABA’ pattern, respectively, with ‘A’ indicating the presumed ancestral state (i.e. that seen in the outgroup) and ‘B’ indicating the derived state.

If there are multiple sequences representing each population, the value for each site can be a proportion, computed from the frequencies of the derived allele in each population:

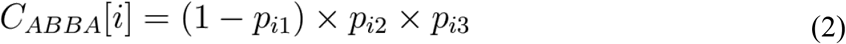

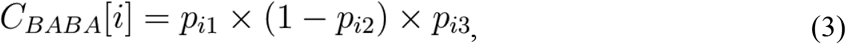

where *p*_*ij*_ is the frequency of the derived allele at site *i* in population *j*. This assumes that the outgroup is fixed for the ancestral state, which is reasonable provided it is sufficiently anciently diverged that few segregating polymorphisms in the ingroups date to before their divergence from the outgroup. Violation of this assumption will result in statistical noise, but as long as the number of sites affected is small, the impact on *D*_FS_ will tend to be modest.

Given equal sample sizes of *n* haploid genotypes per population for both P1 and P2, *D*_FS_ represents the set of partitioned *D* values {*D*_1_, *D*_2_,…,*D*_*k*_,…*D*_*n*_}, in which each value represents a *D* statistic computed using a subset of sites. Specifically, *D*_*k*_ is computed using only ‘ABBA’ sites at which the derived allele occurs *k* times in P2, and ‘BABA’ sites at which the derived allele occurs *k* times in P1. Thus

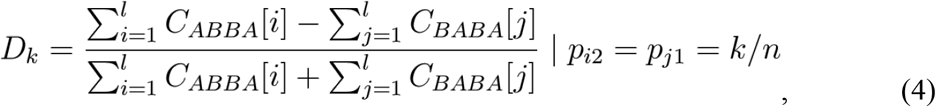

where *p*_*i*2_ is the derived allele frequency in P2 at site *i*, and *p*_*j*1_ is the derived allele frequency in P1 at site *j*.

Finally, since different numbers of sites will contribute to each entry, and each site contributes differently to overall *D* depending on the allele frequencies in each population, each of the partitioned *D* values making up *D*_FS_ is assigned a weighting, 0 ≤ *w*_*k*_ ≤ 1, representing its proportional contribution to overall *D*:

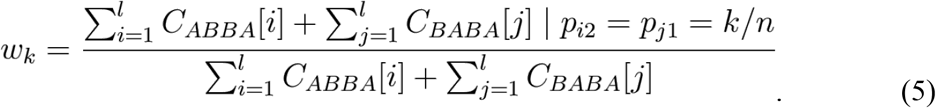

*D*_FS_ can be computed from a joint site frequency spectrum (SFS). This can be a polarised (‘unfolded’) 3-dimensional SFS (i.e. giving the frequency of the derived allele for three populations), or an unpolarised 4-dimensional SFS, in which the outgroup can be used for polarization. We provide code for computation of *D*_FS_ from an input SFS at https://github.com/simonhmartin/dfs.

### Simulations

In order to explore the behaviour of *D*_FS_ over a wide range of parameters, we performed simulations of the 3D SFS using *moments* (Jouganous et al. 2017), which is based on a moment representation of the diffusion equation. Because the simulated SFS is polarised, we did not simulate an outgroup population. This emulates empirical studies in which a suitable outgroup is available for identification of the ancestral allele (as described below). Custom scripts for running single or batch simulations are provided at https://github.com/simonhmartin/dfs.

### Analysis of empirical data

We analysed six previously published whole genome data sets from plants and animals. Due to the different qualities and sequencing depths of each data set, different filtering criteria were used, but otherwise the analysis pipeline was consistent.

*Heliconius* butterfly genotype data (Martin et al. 2019) were accessed from https://doi.org/10.5061/dryad.sk2pd88. Several combinations of populations allow for tests for introgression between *Heliconius melpomene* and either *H. timareta* or *H. cydno*. Frequencies were polarised using the outgroup *H. numata*.

*Arabidopsis* genotype data (Marburger et al. 2019) were obtained from the authors. Only genotypes supported by an individual read depth ≥ 5 were considered, and sites genotyped in fewer than 80% of individuals were excluded. We tested for introgression from tetraploid *Arabidopsis arenosa* (populations BGS, BRD, GUL, SEN, TRE) into tetraploid *A. lyrata* (populations KAG. LIC, MAU, MOD, SCB, SWA). For P1 (the putative non-admixed population), we used diploid *A. lyrata* (populations PEQ, PER, VLH). Frequencies were polarised using the outgroup *H. halleri*.

*Phoenix* date palm genotype data (Flowers et al. 2019) were retrieved from https://doi.org/10.5061/dryad.tm40gd8. Only genotypes supported by an individual read depth ≥ 8 were considered. Following the original study, we tested for introgression from the wild *Phoenix theophrasti* into cultivated North African *P. dactylifera* (represented by samples from Morocco), using middle-eastern *P. dactylifera* (represented by samples from Iraq) as P1. Putatively admixed individuals of the donor *Phoenix theophrasti* were excluded. Frequencies were polarised using the outgroup *P. canariensis*.

American sparrow genotype data (Walsh et al. 2018) were retrieved from https://doi.org/10.5061/dryad.gt12rn3. Only genotypes supported by an individual read depth ≥ 4 were considered. We tested for introgression between sympatric populations of Saltmarsh sparrow *Ammospiza caudacuta* and the recent colonist Nelson’s sparrow *A. nelsoni*. Allopatric A. *nelsoni* was used as P1. Frequencies were polarised using the outgroup seaside sparrow *A. maritima*.

Stickleback genotype data (Dixon et al. 2019) were obtained from the authors. Only genotypes supported by an individual read depth ≥ 5 were considered. We tested for introgression between the ninespine stickleback *Pungitius pungitius* and the Armur stickleback *P. sinensis*. Since no allopatric population of *P. sinensis* was available, we used the closely related Sakhalin stickleback *P. tymensis* as P1. Chromosome 12 was excluded due to its history of suppressed recombination in *P. pungitius*. Frequencies were polarised using the outgroup threespine stickleback *Gasterostreus aculeatus*.

A hominid genotype data set was generated by combining human data (The 1000 Genomes Project Consortium 2015) (phase 3 data set retrieved from ftp.1000genomes.ebi.ac.uk/vol1/ftp/release/20130502/) with that for an Altai Neanderthal (Prüfer et al. 2014) retrieved from http://cdna.eva.mpg.de/neandertal/altai/AltaiNeandertal/. The latter was filtered to remove SNPs at CpG sites (as annotated by the authors), with a genotype quality < 30, and with a read depth < 10 or > 150. As an outgroup to polarise frequencies, we added the chimpanzee allele based on an alignment of syntenic regions retrieved from http://hgdownload.cse.ucsc.edu/goldenpath/hg19/vsPanTro6/. Only chromosome 1 was considered for the hominid analysis to reduce processing time.

In all cases, allele frequencies and polarised site frequency spectra (SFS) were computed using the scripts freq.py and sfs.py, available at https://github.com/simonhmartin/genomics_general.

## Data Accessibility

All empirical data analysed were based on previously published data sets. Processed genotype files, as well as and plotted results underlying all figures are available from the Zenodo digital repository (DOI: XXX).

## ACKNOWLEDGEMENTS

We thank Steven Van Belleghem, Konrad Lohse and Alex Twyford for helpful comments on the manuscript. SHM was funded by a Royal Society University Research Fellowship (Grant number URF\R1\180682).

