## Supplementary material for "Signatures of introgression across the allele frequency spectrum": Figures S1-S5

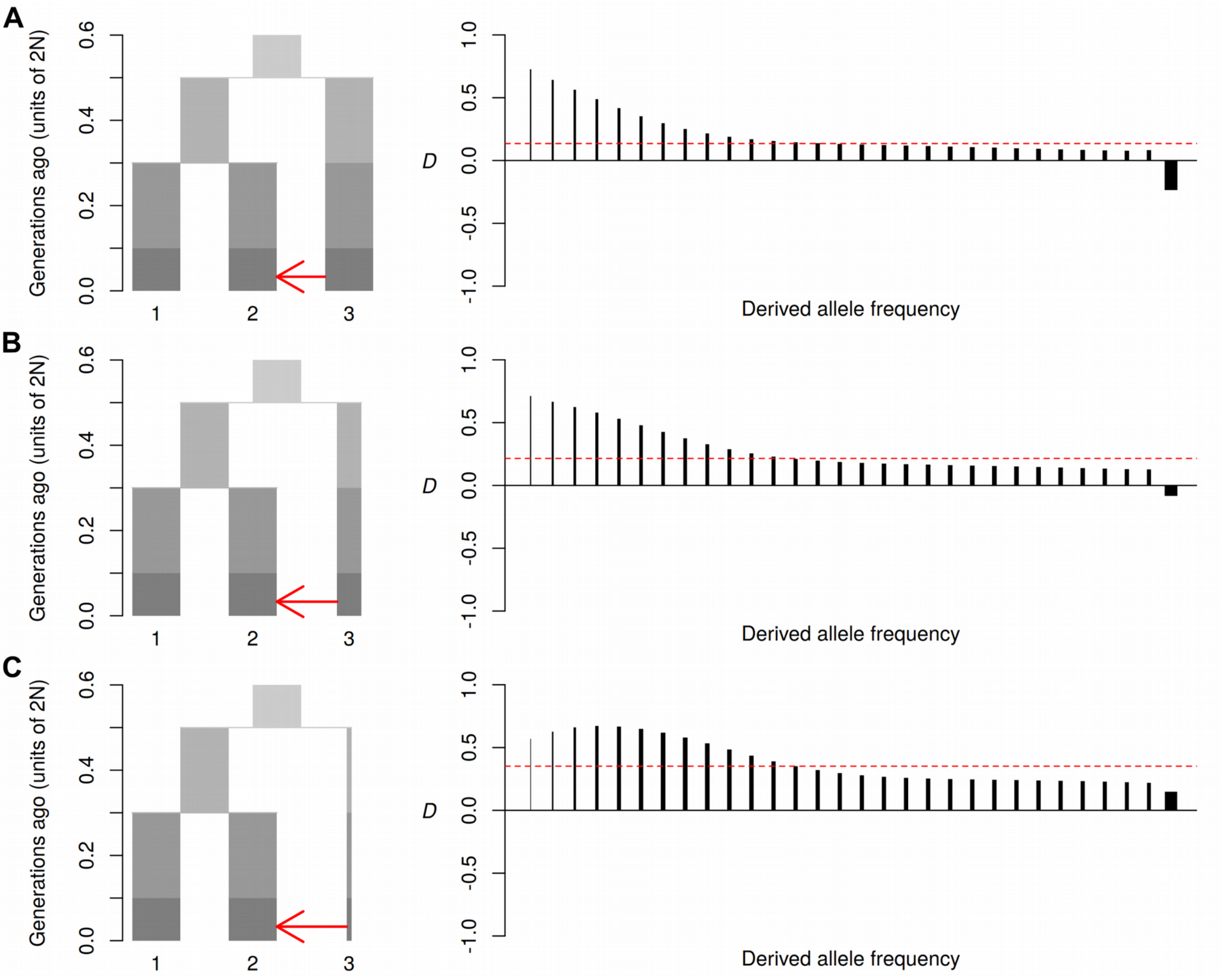


**Figure S1. Lower population size in the donor population alters $D_{FS}$.** Diagrams on the left show the simulated models. Plots on the right show $D_{FS}$ (See Figure 2 in the main paper for full details). Reducing the population size of the donor population (P3) to 50% (panel B) or 10% (panel C) has three effects. Firstly, it causes the overall *D* value to increase. This occurs because the small size of P3 will have allowed many derived alleles to drift to fixation, meaning that introgression will tend to leave a stronger signal of shared derived alleles. Secondly, there can be a rounding of $D_{FS}$ (see panel C) if P3 is very small and migration is high. This also reflects the large number of fixed derived alleles in P3, which will tend to occur immediately at intermediate frequency in the recipient population (P2) following strong gene flow. Finally, it tends to remove negative $D_{FS}$ values in the highest frequency bin (compare the right-most bars in panels A and C). These negative values reflect derived alleles that had fixed in both P1 and P2 but are still segregating in P3 due to ILS. At these sites, gene flow from P3 to P2 has the counter-intuitive effect of reducing the number of fixed derived alleles in P2 that are shared with P3. P1 is unaffected by introgression and therefore retains its fixed derived alleles, creating the imbalance in the high-frequency bin. By decreased the effective population size of the donor population P3, we reduce segregating derived alleles due to ILS, and thereby also reduce the inverted signal. Migration rate ($2Nm$) = 2.

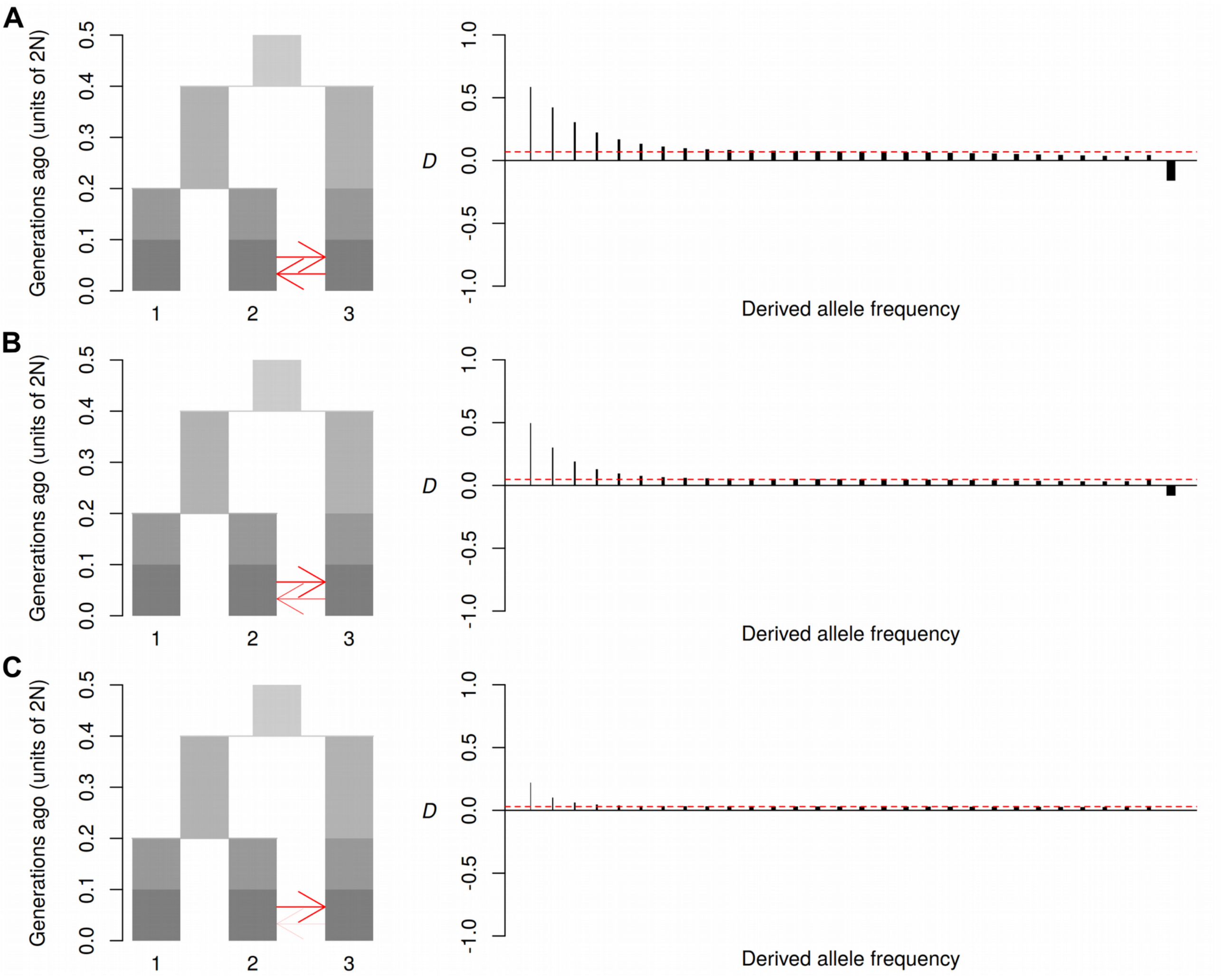


**Figure S2. Relaxing assumptions about the directionality of introgression.** Diagrams on the left show the simulated models. Plots on the right show $D_{FS}$ (See Figure 2 in the main paper for full details). Here we investigate the influence of violation of model assumptions with regard to the direction in introgression. Panel A tests a model in which intogression between P2 and P3 is completely bi-directional ($2Nm = 1$). This shows that the signature of excess allele sharing among low frequency bins is retained and similar to that under unidirectional introgression into P2 (Figure 2A). Panels B and C show reductions in the extent of gene flow into P2 from P3 to 50% (panel B) and 10% (panel C) of that from P2 into P3. In both cases the signal is reduced, but the excess among low-frequency bins is not eliminated. This shows that even strong imbalances in the direction of recent introgression should not necessarily eliminate the expected peak among low frequencies.

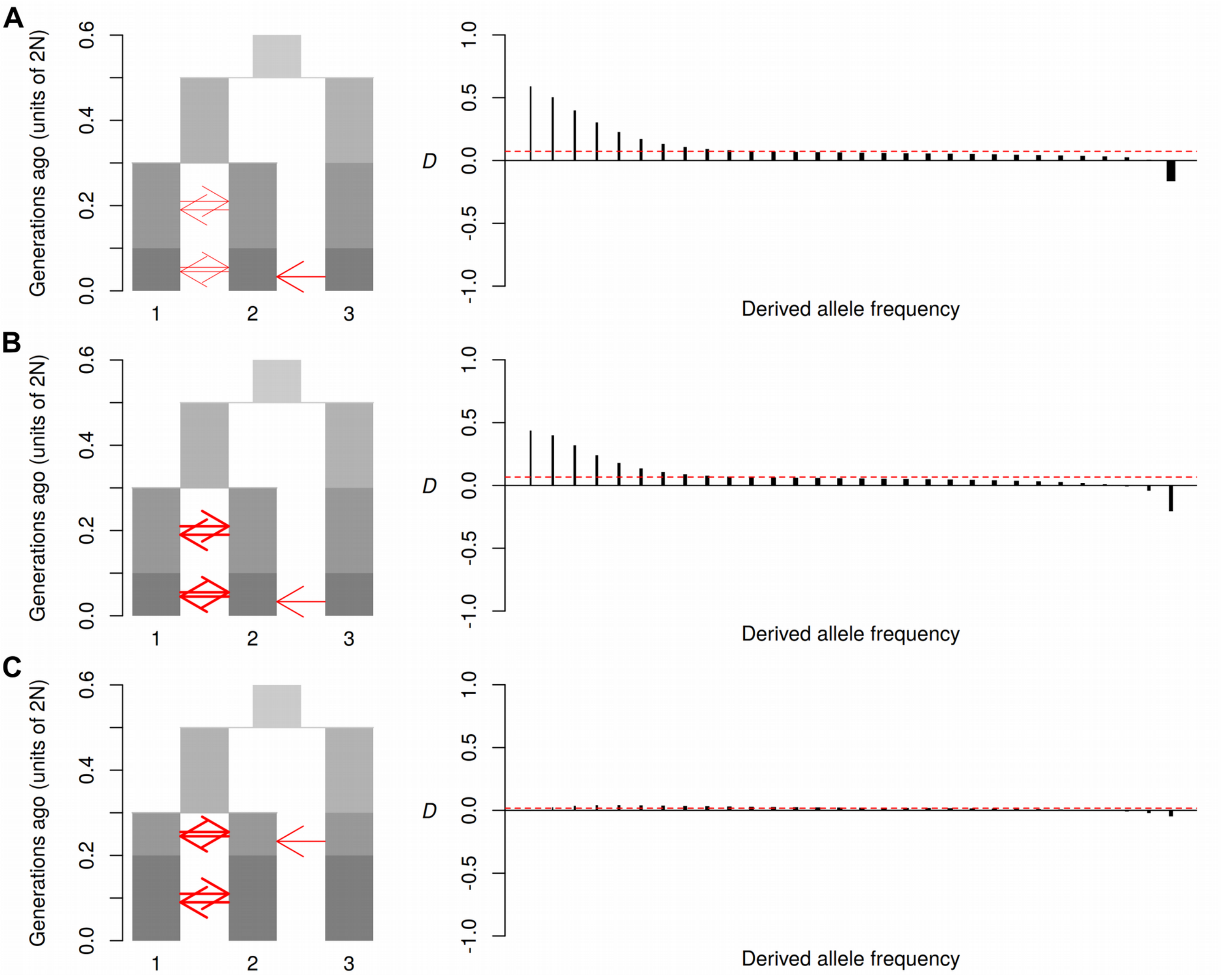


**Figure S3. Relaxing the assumption of isolation between P1 and P2.** Diagrams on the left show the simulated models. Plots on the right show $D_{FS}$ (See Figure 2 in the main paper for full details). Here we investigate the influence of violation of model assumptions with regard to the level of isolation between the focal populations P1 and P2. Panel A tests a model in which continuous migration between P1 and P2 occurs at a rate half that of the level of recent introgression from P3 into P2 ($2Nm$ = 0.5 vs 1). This has minimal impact on the shape of $D_{FS}$ compared to the standard model (Figure 2A). Panel B tests a model in which continuous migration between P1 and P2 occurs at a rate double that of the level of recent introgression from P3 into P2 ($2Nm$ = 2 vs 1). This has a detectable impact on the shape of $D_{FS}$, but the characteristic peak at low frequencies is retained. Panel C tests a model in which introgression occurs further in the past, and continuous migration between P1 and P2 occurs at a rate double that of the historical introgression event. This causes nearly complete erosion of the signal of the older introgression event, especially among low-frequency alleles.

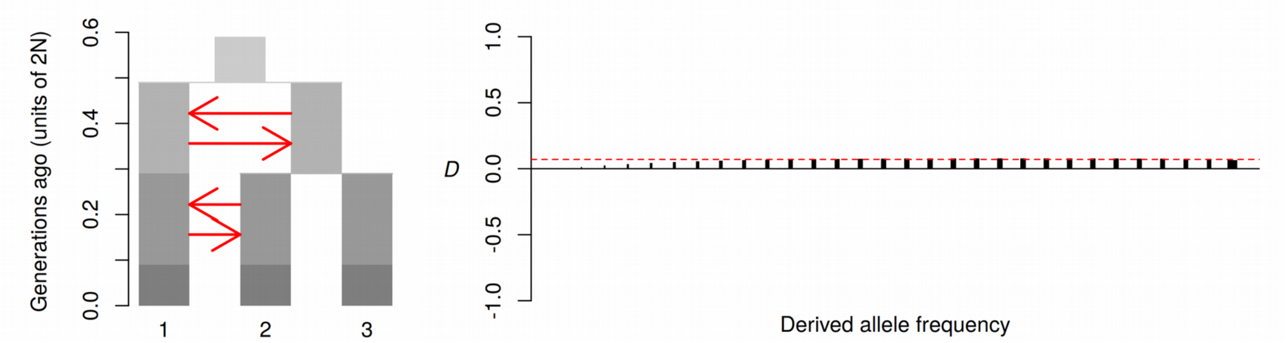


**Figure S4. The effect of ancestral structure on $D_{FS}$.** The diagrams on the left show the simulated model. This model simulates a possible violation of the assumptions of the ABBA BABA test. Whereas P3 is conventionally a sister to both P1 and P2, here the ancestral population is structured with ongoing migration ($2Nm = 2$), and P3 branches off in a 'budding' process, such that it is expected to share more variation with P2 than P1. Thus, a non-zero *D* statistic may occur in the absence of introgression (Eriksson and Manica 2012, Yang et al. 2012). $D_{FS}$ shown on the right indicates that the excess allele sharing is restricted to higher frequencies and is absent from the lowest frequency bins. This is similar to the result of Yang et al. (2012), and indicates that a scenario of ancestral structure may be distinguishable from true recent introgression.

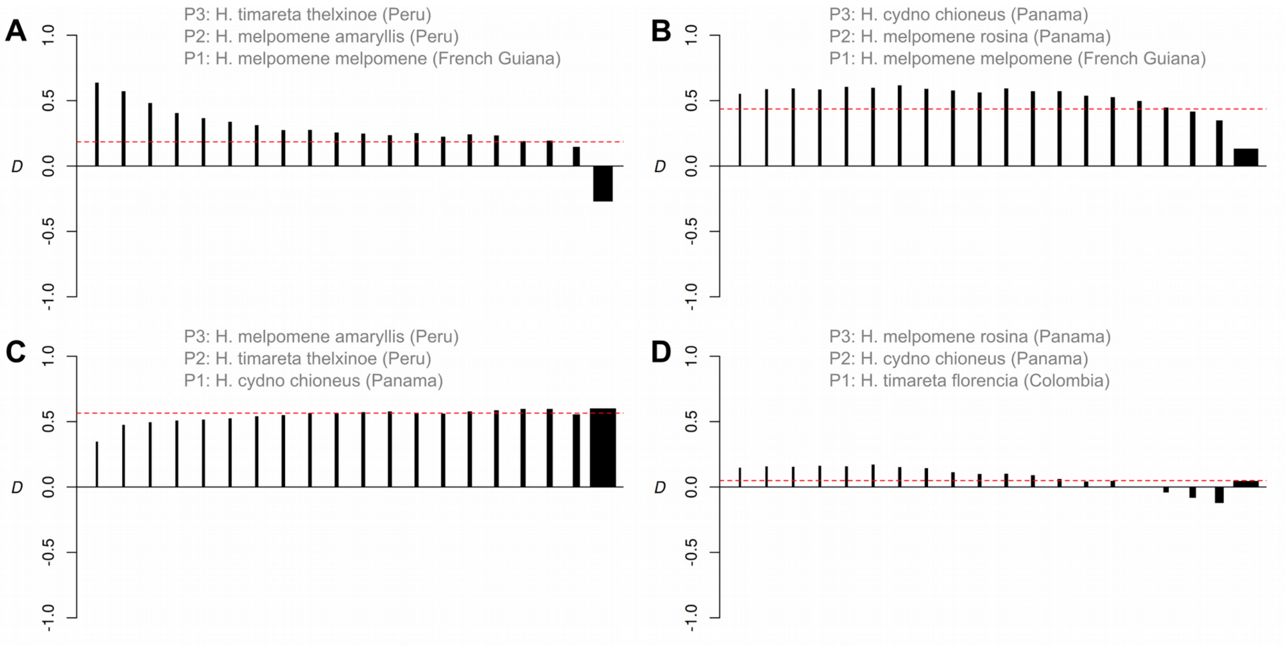


**Figure S5. Comparison of $D_{FS}$ using four different sets of *Heliconius* populations. A.** Introgression from *H. timareta thelxinoe* into *H. melpomene amaryllis* in Peru, using the allopatric *H. melpoemene melpomene* from French Guiana as P1. This detects a pattern consistent with recent gene flow. **B**. Introgression from *H. cydno chioneus* into *H. melpomene rosina* in Panama, again using the allopatric *H. melpoemene melpomene* from French Guiana as P1. The observed pattern is more consistent with older gene flow. **C.** The same hybridising pair as panel A, except with *H. t. thelxinoe* as the focal population, and *H. c. chioneus* as P1. The dip at low frequencies is most likely caused by the fact that *H. c. chioneus* has also received introgressed variation through interbreeding with *H. m. rosina* in Panama. This highlights the importance of using an allopatric P1 population. **D.** The same hybridising pair as B, except with *H. cydno chioneus* as the focal population and a different *H. timareta* population as P1. The weaker signal might result from lower rates of gene flow into *H. cydno*, but the fact that *H. tinmareta* is not allopatric from H. melpomene also makes this result difficult to interpret.
